# Structure of the Monkeypox profilin-like protein A42R reveals potential function differences from cellular profilins

**DOI:** 10.1101/2022.08.07.503103

**Authors:** George Minasov, Nicole L. Inniss, Ludmilla Shuvalova, Wayne F. Anderson, Karla J. F. Satchell

**Affiliations:** Department of Microbiology-Immunology, Northwestern University Feinberg School of Medicine, Chicago, Illinois, USA 60611; Center for Structural Genomics of Infectious Diseases, Northwestern University Feinberg School of Medicine, Chicago, Illinois, USA 60611; Department of Pharmacology, Northwestern University Feinberg School of Medicine, Chicago, Illinois, USA 60611; Department of Biochemistry and Molecular Genetics, Northwestern University Feinberg School of Medicine, Chicago, Illinois, USA 60611

## Abstract

The infectious disease human monkeypox is spreading rapidly in 2022 causing a global health crisis. The genomics of the monkeypox virus (MPXV) have been extensively analyzed and reported, although little is known about the virus-encoded proteome. In particular, there is almost no structure information about MPXV proteins, other than computational models. Here we report a 1.52 Å x-ray structure of the MPXV protein A42R, the only MPXV-encoded protein with a known structure. This protein shows sequence similarity to profilins, which are cellular proteins known to function in regulation of actin cytoskeletal assembly. However, structural comparison of the A42R with known members of the profilin family reveals critical differences that support prior biochemical findings that this protein only weakly binds actin and does not bind poly(L-proline). In addition, the analysis suggests that this protein may have distinct interactions with phosphatidylinositol lipids. Overall, our data suggest that the role of A42R in replication of orthopoxviruses may not be easily determined by comparison to cellular profilins. Further, our findings support a need for increased efforts to determine high-resolution structures of other MPXV proteins to inform physiological studies of the poxvirus infection cycle and to reveal potential new strategies to combat human monkeypox should the infection become more common in the future.

## INTRODUCTION

Monkeypox virus (MPXV) is a poxvirus in the *Orthopoxvirus* genus. The virus is closely related to other human pathogens, including variola major virus (VARV), which causes smallpox, cowpox virus (CPXV), and vaccinia virus (VACV).^1^ MPXV was first described in 1958 during an outbreak among macaques originating from Singapore that were imported into Denmark for polio-vaccine research.^2^ The disease human monkeypox is typified by headache, fever, and flu-like symptoms with characteristic pox lesions appearing shortly after symptom onset.^3,4^ Although most patients resolve the infection in two to four weeks, the disease can be fatal with a case-fatality ratio of 10.6% for infections by MPXV Clade 1 strains (formerly known as the Central African clade), and 3.6% for infections by MPXV Clade 2 and 3 strains (formerly known as the West African Clade).^5^

Human monkeypox is most commonly a zoonotic infection contracted from wild animals and then passed human-to-human by direct contact.^5^ The disease primarily occurs in the rain forest areas of Western and Central Africa, where discrete outbreaks occur impacting only a small number of patients, except in the Democratic Republic of Congo, which has reported more than 1000 cases per year since 2005.^4,5^ The previously largest outbreak outside of Africa occurred in 2003 when 35 cases in the Midwestern United States were linked to contact with prairie dogs from a distributor that also imported African rodents.^6^ The now largest on-going outbreak of MPXV began in April 2022 and is linked to more than 28,000 cases across 88 countries, predominantly in the United States, Europe, and Brazil.^7^ The World Health Organization has declared human monkeypox a public health emergency of global concern.^8^

MPXV is an enveloped, brick-shaped virus with a linear, double-stranded DNA genome. The representative MPXV genome from isolate Zaire-96-I-16 is 196.8 kB with 191 open reading frames and derives from Clade 1.^9^ The 2022 outbreak is a new branch on the phylogenetic tree within Clade 3 and is most similar to the strain that caused a 2017-2018 outbreak in Nigeria that spread to the United Kingdom.^10^ There are already more than 400 genomes of the 2022 outbreak strain available at National Center for Bioinformatic Information (NCBI) GenBank with genomes of 184.7 – 198.0 kb annotated as having from 143-214 open reading frames. Analysis of the genomes reveals the strain derives from a West African isolate and differs from 2018-2019 isolates by 50 single nucleotide polymorphisms.^10^

Despite extensive knowledge on genomics and epidemiology of MPXV, the proteome of MPXV is not well studied. Further, there are almost no protein structures encoded from the MPXV genome other than computer models. The A42R protein that is encoded by the MPXV gp153 locus has significant amino acid sequence identity to eukaryotic cell profilin proteins^11^ (Figure 1A). Proteins from the profilin family are actin-binding proteins that participate in F-actin assembly and regulation.^12-14^ The VACV profilin-homology protein (VACV A42R), which is 98% identical to the MPXV A42R (Figure 1A), is a late expressing viral protein^15^ with weak affinity for actin (19). A peptide of VACV A42R (^88^YAPVSPIVI^96^) is a CD8+ T cell epitope.^16^ Genetic knockout of the open reading frames for the profilin-homology proteins of VACV and CPXV viruses demonstrated these proteins are not essential for replication of poxviruses *in vitro*.^15,17^ Thus, the role of the highly conserved profilin-like proteins in the poxvirus infection cycle is not known. Here, we report the structure of A42R, the first structure of a MPXV protein deposited to the RCSB Protein Data Bank (PDB). Although the structure detailed here was first deposited and released back in 2014, it remains the only protein from MPXV in the PDB and the only poxvirus profilin-like protein with a known structure. Comparison with structures of human and bovine profilin proteins reveals significant differences in functional regions, indicating that the role of MPXV A42R in the viral life cycle may not be easily determined based solely on our understanding of cellular profilin function.

**FIGURE 1.**
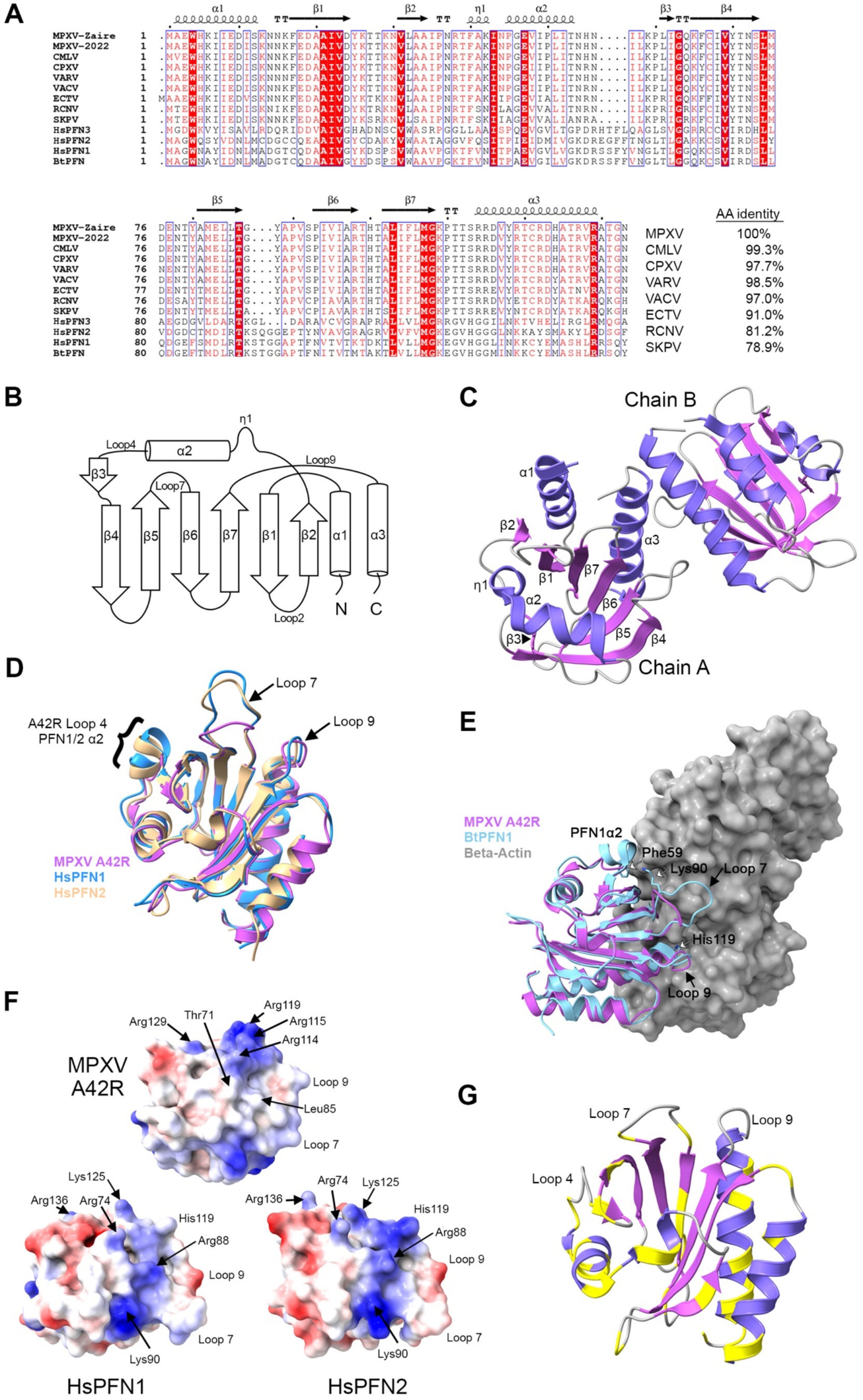
Structural analysis of MPXV A42R. (A) Sequence alignment of MPXV A42R with orthopoxvirus profilin-like proteins and mammalian profilin proteins. (B) Schematic of the β-strand and α-helical arrangement of the A42R protein. (C) Α cartoon representation of the x-ray structure of MPXV-Zaire A42R structure (PDB code 4QWO) with β-strands in violet and α-helices in dark purple. Strands and helices in Chain A are marked as indicated in panel B. (D) Overlay of MPXV A42R (violet) with *Homo sapiens* (human) *Hs*PFN1 (PDB code 1FIL, blue, unpublished) and human *Hs*PFN2 (PDB code 1D1J, beige, ref. ^20^). Loops are labeled according to MPXV A42R structure as shown in Panel B. (E) Overlay of MPXV A42R (violet) with structure of Bos taurus (bovine) *Bt*PFN1 (light blue) bound to beta-actin (grey) (PDB code 3UB5, ref. ^23^). Loops are labeled according to MPXV A42R structure as shown in Panel B. (F) Electrostatic surface projections of MPXV A42R, HsPFN1, and HsPFN2 (PDB codes 4QWO, 1FIL, and 1D1J, respectively). Negatively charged residues are red and positively charged residues are blue with residues noted in text indicated. Loops as indicated in Panel E are shown for orientation of the actin binding region. (G) Structure of MPXV A42R oriented as in Panel D and colored as in Panel C with residues that vary in any of the nine orthopoxviruses in Panel A alignment indicated in yellow. Loops as marked as in Panel B.

## RESULTS

### MPXV A42R resembles human profilins

The open reading frame encoding MPXV A42R was amplified and cloned from MPXV Zaire-96-I-16 strain DNA (NCBI accession NP_536579.1, ref. ^9^). The A42R protein was expressed in *E. coli* in selenomethionine-containing media and purified as described in the methods. The protein was then crystallized in a 25% (w/v) PEG 1500 and 0.1M MMT buffer solution at pH 9.0. The diffraction quality crystals belonged to the P2_1_2_1_2_1_ space group. The structure was solved using single-wavelength anomalous dispersion (SAD) method and refined to 1.52 Å resolution.

The asymmetric unit consisted of two polypeptide chains. Full length Chain A was intact across all 133 residues of A42R and also contained an N-terminal alanine residue originating from the tobacco-etch mosaic virus (TEV) protease recognition site. Chain B contained only residues 2-133. The overall structure formed a seven-stranded anti-parallel beta sheet surrounded by three alpha helices and one partial helix (Figure 1 B,C). A Dali search^18^ for protein structures with similar fold revealed that MPXV A42R closely resemble structures determined for the ubiquitously expressed human profilin-1 (PFN1, 32% identity), and the neuronal specific profilin-2 (PFN2, 26% identity), which share a well-conserved fold.^12^ There is also weaker structural homology to *Arabidopsis thaliana* profilin-3 (PFN3, 23% identity), which also functions in actin dynamics.^19^ Therefore, our structure represents a family of profilin-like orthopoxviral proteins that may play a role in altering actin dynamics during viral replication in cells.

When A42R was overlaid with human PFN1 (PDB code 1FIL, unpublished) and PFN2 (PDB code 1D1J, ref. ^20^), the backbones of the structures aligned with a root mean-square deviation (RMSD) of 0.8 Å (117 pruned Cα pairs) with PFN1 and an RMSD of 1.0 Å (110 pruned Cα pairs) with PFN2. The major differences between structures are located in the loop regions (Figure 1D). In particular, the loop 4 between α2 and β2 in A42R from MPXV is four residues shorter than the same loop in PFN1 and PFN2. In the human profilin proteins, part of this loop forms a helix (PFN1/PFN2 α3). Thus, MPXV A42R has only three helices surrounding the seven-strand beta-sheet compared to the four helices that typifies other profilin proteins. In addition, loop 7 between strands β5 and β6 has a three-residue deletion that makes this loop shorter. Interestingly, this loop in human PFN3 is also shorter than in PFN1 and PFN2 (Figure 1A), although the impact of this on function is not known. Structural analysis of the computational model of the A42R homolog from ectromelia virus (ECTV) also predicted significant differences in these loops compared to human profilins.^21^

### MPXV A42R structure analysis supports that poxvirus profilin-like proteins weakly bind to actin

Profilins interact directly with actin to sequester actin monomers and then deliver the ATP-actin complex to the growing end of an actin filament.^14^ Biochemical studies have shown that VACV A42R binds actin, but the binding affinity is weak compared to both PFN1 and PFN2.^22^ Comparative analysis of the x-ray structure of MPXV A42R with mammalian PFN1 and PFN2 suggest that shortening of these loop regions could have significant functional impact on the binding affinity of MPXV A42R to actin. Overlay of the structure of A42R from MPXV with the structure of *Bos taurus* (bovine) PFN1 (*Bt*PFN1) with bound beta-actin (PDB code 3UB5, ref. ^23^) shows that loops 4, 7, and 9 participate in formation of the patch on the surface that interacts with the surface of the actin (Figure 1E). However, the residues involved in this critical interaction are different in A42R. In particular, amino acid differences in A42R loop 9 eliminated the critical interaction of PFN1 His119 with a hydrophilic pocket of the beta-actin formed in part by Tyr133 and Tyr169. (Figure 1E). Loss of this histidine interaction with actin may in part be replaced by a weak hydrogen bond interactions between Arg115 and Arg119 of MXPV A42R with Glu61 of actin. Further, Phe59 from α3 of PFN1 forms a stacking interaction with His173 of actin. This critical Phe residue is lost in the replacement of *Bt*PFN1 α3 with the shorter loop 4 in MPXV A42R. Finally, Lys90 from the loop 7 of PFN1 interacts with Asp288 of the actin, but this residue is missing in MPXV A42R due to the deletion in the loop 7.

### PIP2 interaction surface is distinct in MPXV A42R compared to profilins

Profilins are known to regulate actin polymerization through interaction with phosphatidylinositol-(4,5)-bisphosphate (PIP2) at cell membranes. PIP2 binding to profilins also sequesters PIP2 from turnover by phospholipase C-gamma.^14^ PIP2 binding to human profilins is proposed to require a positively charged surface that involves residues Arg74, Arg88, Lys90, and Lys125 (Figure 1F) and that partially overlaps the actin-binding face.^12,24^ Comparison of electrostatic projections of the proposed PIP2 binding surface shows MPXV A42R has significant reorganization of basic and hydrophobic regions compared to PFN1 and PFN2 (Figure 1F). In particular, mutagenesis studies demonstrated PFN1 Arg88 and surrounding residues are particularly critical for PIP2 binding.^12,25^ In MPXV A42R, PFN1 Arg88 is replaced by Leu85 and nearby Lys90 of PFN1 is lost because of deletion in the loop 7. Further, Arg74 in PFN1 is changed to a threonine (Thr71) in MPXV A42R. Thus, the surface in this region is hydrophobic rather than basic. Biochemical studies have nonetheless shown that MPXV A42R does bind PIP2.^26^ Structural analysis of the MPXV A42R suggests that PIP2 may bind via an arginine patch comprised of Arg114, Arg115, and Arg119. This suggestion is consistent with a role of the C-terminal alpha-helix of PFN1, and specifically Lys125 (same position as MPXV A42R Arg119) as a contributor to PIP2 binding.^12,25^ It has also been suggested that in PFN1 Arg136 contributes to a second PIP2 binding site.^27^ This residue corresponds to Arg129 in MPXV A42R and interestingly is conserved in all profilin and viral profilin-like proteins (Figure 1A), indicating that this PIP2 binding site may be conserved in the viral profilin-like proteins.

### Possible interactions of MPXV A42R with other proteins

In addition to interaction with actin and PIP2, profilins bind to actin-associated proteins and other cellular proteins.^13,14,28^ Many of these interactions are through the poly(L-proline) interaction region formed by the N- and C-terminal helices and the binding of proteins with proline repeat domains is regulated by phosphorylation of a serine located on the C-terminal helix.^14^ However, VACV A42R does not bind poly(L-proline)^22^ although the poxvirus profilin-like proteins do have a threonine (Thr131) at the position of the phosphorylated serine. Thus, it is unlikely that MPXV A42R binds to proteins with proline repeat domains despite the presence of a Trp4 (Trp5 in VACV A42R) known to disrupt binding to poly(L-proline) proteins when changed by site-directed mutagenesis.^28^

PFN1 is also known to interact with microtubules through Met114 and Gly118.^29^ As noted above, in MPXV A42R, these residues are changed to form an arginine patch and we posit this patch may contribute to PIP2 binding. Since the surface at this face is dramatically changed in MPXV A42R, we speculate that this protein may not bind microtubules.

Finally, the ECTV A42R was shown to bind and co-localize with cellular tropomyosin, a protein that polymerizes along actin filaments to regulate binding of actin-associated proteins to the filaments.^21^ Tropomyosin has not been recognized as ligand for human profilins.^13,14^ Although the regions of the ECTV A42R protein that bind tropomyosin are not yet known, this finding indicates that poxvirus profilin-like proteins may engage in unique protein-protein interactions and regulate actin in a manner distinct from human profilins.

### A42R structure is highly conserved across the Orthopoxviruses

To understand if our structure here would impact our understanding of other poxviruses, we compared the sequence of MPXV A42R with other poxvirus profilin-like proteins. MPXV A42R differs from VARV and VACV by only two amino acids and from CPXV by three amino acids (Figure 1A). The sequence alignment with other more distant orthopoxviruses also show high identity across all residues with the Skunkpox virus (SKPV) being the most distant homolog with 79% identity. In total, we identified 39 single residue differences across nine homologous orthopoxvirus A42R proteins and mapped them on the surface of our structure (Figure 1G). These differences span along the entire protein and are, in general, modest substitutions with retention of residue characteristics. Interestingly, residues 17-24 that form the β1 strand are 100% identical across all nine homologs suggesting a key role for this region in protein stability and/or function. All orthopoxvirus profilin-like proteins also have loops 4 and 7 that are shorter compared to cellular profilins. Further, the presence of all three arginine residues that form the arginine patch in MPXV is also conserved across the orthopoxviral proteins, except for SKPV which has Arg119 substituted by an asparagine residue. The conservation of the potential functional elements suggests that structure of MPXV A42R is a valid homologous representative across the orthopoxvirus genus.

## DISCUSSION

Overall, our structure of MPXV A42R is not inconsistent with the hypothesis that this protein could function to regulate actin remodeling, similar to the host profilins. However, closer structural comparisons reveal that A42R has significant changes that may impact its interactions with other proteins or ligands and that these differences could indicate distinct functions of this protein in cells. As most biochemical studies of A42R were conducted in the 1990s, our work highlights the value in further investigation of the function of A42R and other viral profilin-like proteins in the viral life cycle. New molecular and biochemical studies are needed to determine biologically relevant binding partners of A42R and the role of these interactions during viral replication in cells.

Although the VACV and CPXV profilin-like proteins were not essential for viral replication in cultured cells,^15,17^ it remains possible that this protein is important during infection in different cell types or in a clinical setting. Neither the VACV nor CPXV strains with the open reading frames for the profilin-like proteins interrupted were tested in animals and thus the impact of the protein on viral replication *in vivo* remains unknown. Further, our analysis of the first structurally characterized protein from MPXV suggests that there would be a value in conducting structural studies of other MPXV proteins, even those that may be similar to host proteins with known structure. Indeed, it may be critical to reveal more about the physiology of MPXV if the current outbreak continues to spread or if outbreaks of similar scope begin to recur more frequently. Such efforts could inform development of new therapeutics. Currently, only tecovirimat, an envelope protein wrapping inhibitor, and brincidofovir, a nucleoside analog, are FDA-approved for treatment of monkeypox.^30^ Structural modeling may serve to fill a knowledge gap in the absence of x-ray structures. Indeed, a recent preprint reported analysis of the genome of the newly circulating strain of MPXV and predicted the 3D structures of the protein encoded by 123 different open reading frames.^31^ However, for structure-guided drug discovery and establishing structure-activity relationships, high resolution crystal structures such as the 1.52 Å resolution reported here are still of high value to conduct advanced biological and translational studies of this understudied pathogen.

## MATERIALS AND METHODS

### Protein expression, purification, and crystallization

The open reading frame encoding A42R was amplified from genomic DNA of the MPXV Zaire-96-I-16 strain (NCBI accession NC_003310.1) into vector pMCSG53.^32^ Recombinant A42R was expressed and purified from *E. coli* BL21(DE3)(Magic) cells according to published methods.^33^ The final purified protein was set up for crystallization at 2.43 mg/ml in 0.5 M sodium chloride, 0.01 M Tris-HCl buffer pH 8.3, 5 mM beta-mercaptoethanol as 2 μl crystallization drops (1 m μl protein: 1 μl reservoir solution) in 96-well crystallization plates. Diffraction quality crystals were obtained from condition 0.1 M MMT buffer (pH 9.0), 25% (w/v) PEG 1500 (QIAGEN PACT screen, condition 42).

### Structure determination and refinement

Diffraction data were collected at the 21-ID-G beamline of the Life Science Collaborative Access Team (LS-CAT) at the Advanced Photon Source, Argonne National Laboratory. The data set was processed and scaled with the HKL3000 suite.^34^ The structure was solved by the single-wavelength anomalous dispersion method (SAD). The initial solution went through several rounds of refinement in REFMAC v. 5.8.0258 ^35^ and manual model corrections using Coot.^36^ The water molecules were generated using ARP/wARP ^37^ and ligands were added to the model manually during visual inspection in Coot. Translation– Libration–Screw (TLS) groups were created by the TLSMD server (http://skuldbmsc.washington.edu/~tlsmd/)^38^ and TLS corrections were applied during the final stages of refinement. Molprobity (http://molprobity.biochem.duke.edu/)^39^ was used for monitoring the quality of the model during refinement and for the final validation of the structure. Structure analysis and figure preparation was done with UCSF ChimeraX.^40^

### Sequence analysis

Sequence analysis was done with sequences from MPXV-Zaire-96-I-16 (NP_536579.1), MPXV-2022 Clade 3 (YP_010377149.1), Camelpox virus (CMLV, Q775N7.1), Cowpox virus (CPXV, NP_619961.1), Variola major virus (VARV, ABF24320.1), Vaccinia virus (VACV, QQ05880.1), Ectromelia virus (ECTV, NP_671660.1), Raccoonpox, YP_009143471.1), Skunkpox (SKPV, YP_00928256.1), human PFN1 (NP_005013.2), human PFN2 (NP_444252.1), human PFN3, (NP001025057.1), and *Bos taurus* (NP_001015592.1). The alignment was prepared using Clustal Omega^41^ and ESPript3.^42^

### Data availability

The final models and diffraction data were deposited to the Protein Data Bank (https://www.rcsb.org/) with the assigned PDB code 4QWO.

**TABLE 1.** Data quality and refinement statistics. Values in parentheses are for the outer shell.

| **PDB Accession Code** | **4qwo** |
| --- | --- |
| **Data Collection** |  |
| Space group | $P2_12_12_1$ |
| Unit cell parameters (Å; °) | $a = 41.62, b = 50.50, c = 119.62;$<br>$\alpha = 90.00, \beta = 90.00, \gamma = 90.00$ |
| Resolution range (Å) | 30.00 - 1.52 (1.55 - 1.50) |
| No. of reflections | 39,686 (1,955) |
| $R_{\text{merge}}$ (%) | 8.7 (54.7) |
| Completeness (%) | 100.0 (100.0) |
| $\langle I/\sigma(I) \rangle$ | 31.4 (3.2) |
| Multiplicity | 7.1 (7.2) |
| Wilson $B$ factor | 14.3 |
| **Refinement** |  |
| Resolution range (Å) | 28.81 - 1.52 (1.56 - 1.52) |
| Completeness (%) | 99.9 (99.0) |
| No. of reflections | 37,630 (2,843) |
| $R_{\text{work}}/R_{\text{free}}$, (%) | 14.1/16.6 (17.0/20.2) |
| Protein chains/atoms | 2/2,302 |
| Ligand/Solvent atoms | 42/478 |
| Mean temperature factor (Å<sup>2</sup>) | 16.3 |
| **Coordinate Deviations** |  |
| R.m.s.d. bonds (Å) | 0.010 |
| R.m.s.d. angles (°) | 1.517 |
| **Ramachandran plot** |  |
| Favored (%) | 97.0 |
| Allowed (%) | 3.0 |
| Outside allowed (%) | 0.0 |

## ACKNOWLEDGEMENTS

We thank I. Dubrovska, K. Flores, S. Grimshaw, and K. Kwon for contributions to protein expression, purification, and crystallization. This research used resources of the Advanced Photon Source, a U.S. Department of Energy (DOE) Office of Science User Facility operated for the DOE Office of Science by Argonne National Laboratory under Contract No. DE-AC02-06CH11357. Use of the LS-CAT Sector 21 was supported by the Michigan Economic Development Corporation and the Michigan Technology Tri-Corridor (Grant 085P1000817). This project has been funded in whole or in part with Federal funds from the National Institute of Allergy and Infectious Diseases, National Institutes of Health, Department of Health and Human Services, under Contracts No. HHSN272201200026C (to W.F.A) and HHSN272201700060C (to K.J.F.S).

## REFERENCES

1. Petersen E, Kantele A, Koopmans M et al. (2019) Human Monkeypox: Epidemiologic and Clinical Characteristics, Diagnosis, and Prevention. Infect Dis Clin North Am 33:1027–1043. PMID: 30981594 {Medline}

2. Parker S, Buller RM (2013) A review of experimental and natural infections of animals with monkeypox virus between 1958 and 2012. Future Virol 8:129–157. PMID: 23626656 {Medline}

3. McCollum AM, Damon IK (2014) Human monkeypox. Clin Infect Dis 58:260–267. PMID: 24158414 {Medline}

4. Durski KN, McCollum AM, Nakazawa Y et al. (2018) Emergence of Monkeypox - West and Central Africa, 1970-2017. MMWR Morb Mortal Wkly Rep 67:306–310. PMID: 29543790 {Medline}

5. Bunge EM, Hoet B, Chen L et al. (2022) The changing epidemiology of human monkeypox-A potential threat? A systematic review. PLoS Negl Trop Dis 16:e0010141. PMID: 35148313 {Medline}

6. CDC (2003) Multistate outbreak of monkeypox--Illinois, Indiana, and Wisconsin, 2003. MMWR Morb Mortal Wkly Rep 52:537–540. PMID: 12803191 {Medline}

7. CDC. Monkeypox, https://www.cdc.gov/monkeypox: 2022.

8. WHO. WHO Director-General declares the ongoing monkeypox outbreak a Public Health Emergency of International Concern. (2022). https://www.who.int.

9. Shchelkunov SN, Totmenin AV, Babkin IV et al. (2001) Human monkeypox and smallpox viruses: genomic comparison. FEBS Lett 509:66–70. PMID: 11734207 {Medline}

10. Isidro J, Borges V, Pinto M et al. (2022) Phylogenomic characterization and signs of microevolution in the 2022 multi-country outbreak of monkeypox virus. Nat Med. PMID: 35750157 {Medline}

11. Van Vliet K, Mohamed MR, Zhang L et al. (2009) Poxvirus proteomics and virus-host protein interactions. Microbiol Mol Biol Rev 73:730–749. PMID: 19946139 {Medline}

12. Krishnan K, Moens PDJ (2009) Structure and functions of profilins. Biophys Rev 1:71–81. PMID: 28509986 {Medline}

13. Witke W (2004) The role of profilin complexes in cell motility and other cellular processes. Trends Cell Biol 14:461–469. PMID: 15308213 {Medline}

14. Davey RJ, Moens PD (2020) Profilin: many facets of a small protein. Biophys Rev 12:827–849. PMID: 32661903 {Medline}

15. Blasco R, Cole NB, Moss B (1991) Sequence analysis, expression, and deletion of a vaccinia virus gene encoding a homolog of profilin, a eukaryotic actin-binding protein. J Virol 65:4598–4608. PMID: 1870190 {Medline}

16. Moutaftsi M, Peters B, Pasquetto V et al. (2006) A consensus epitope prediction approach identifies the breadth of murine T(CD8+)-cell responses to vaccinia virus. Nat Biotechnol 24:817–819. PMID: 16767078 {Medline}

17. Xu Z, Zikos D, Osterrieder N et al. (2014) Generation of a complete single-gene knockout bacterial artificial chromosome library of cowpox virus and identification of its essential genes. J Virol 88:490–502. PMID: 24155400 {Medline}

18. Holm L (2022) Dali server: structural unification of protein families. Nucleic Acids Res. PMID: 35610055 {Medline}

19. Qiao Z, Sun H, Ng JTY et al. (2019) Structural and computational examination of the Arabidopsis profilin-Poly-P complex reveals mechanistic details in profilin-regulated actin assembly. J Biol Chem 294:18650–18661. PMID: 31653702 {Medline}

20. Nodelman IM, Bowman GD, Lindberg U et al. (1999) X-ray structure determination of human profilin II: A comparative structural analysis of human profilins. J Mol Biol 294:1271–1285. PMID: 10600384 {Medline}

21. Butler-Cole C, Wagner MJ, Da Silva M et al. (2007) An ectromelia virus profilin homolog interacts with cellular tropomyosin and viral A-type inclusion protein. Virol J 4:76. PMID: 17650322 {Medline}

22. Machesky LM, Cole NB, Moss B et al. (1994) Vaccinia virus expresses a novel profilin with a higher affinity for polyphosphoinositides than actin. Biochemistry 33:10815–10824. PMID: 8075084 {Medline}

23. Porta JC, Borgstahl GE (2012) Structural basis for profilin-mediated actin nucleotide exchange. J Mol Biol 418:103–116. PMID: 22366544 {Medline}

24. Fedorov AA, Magnus KA, Graupe MH et al. (1994) X-ray structures of isoforms of the actin-binding protein profilin that differ in their affinity for phosphatidylinositol phosphates. Proc Natl Acad Sci U S A 91:8636–8640. PMID: 8078936 {Medline}

25. Sohn RH, Chen J, Koblan KS et al. (1995) Localization of a binding site for phosphatidylinositol 4,5-bisphosphate on human profilin. J Biol Chem 270:21114–21120. PMID: 7673143 {Medline}

26. Gieselmann R, Kwiatkowski DJ, Janmey PA et al. (1995) Distinct biochemical characteristics of the two human profilin isoforms. Eur J Biochem 229:621–628. PMID: 7758455 {Medline}

27. Lambrechts A, Jonckheere V, Dewitte D et al. (2002) Mutational analysis of human profilin I reveals a second PI(4,5)-P2 binding site neighbouring the poly(L-proline) binding site. BMC Biochem 3:12. PMID: 12052260 {Medline}

28. Kaiser DA, Pollard TD (1996) Characterization of actin and poly-L-proline binding sites of Acanthamoeba profilin with monoclonal antibodies and by mutagenesis. J Mol Biol 256:89–107. PMID: 8609617 {Medline}

29. Henty-Ridilla JL, Juanes MA, Goode BL (2017) Profilin Directly Promotes Microtubule Growth through Residues Mutated in Amyotrophic Lateral Sclerosis. Curr Biol 27:3535–3543 e3534. PMID: 29129529 {Medline}

30. Delaune D, Iseni F (2020) Drug Development against Smallpox: Present and Future. Antimicrob Agents Chemother 64. PMID: 31932370 {Medline}

31. Parigger L, Krassnigg A, Grabuschnig S et al. (2022) Preliminary structural proteome of the monkeypox virus causing a multi-country outbreak in May 2022. Research Square.

32. Stols L, Gu M, Dieckman L et al. (2002) A new vector for high-throughput, ligation-independent cloning encoding a tobacco etch virus protease cleavage site. Protein Expr Purif 25:8–15. PMID: 12071693 {Medline}

33. Shuvalova L (2014) Parallel protein purification. Methods Mol Biol 1140:137–143. PMID: 24590714 {Medline}

34. Minor W, Cymborowski M, Otwinowski Z et al. (2006) HKL-3000: the integration of data reduction and structure solution - from diffraction images to an initial model in minutes. Acta Crystallographica Section D 62:859–866.

35. Murshudov GN, Skubak P, Lebedev AA et al. (2011) REFMAC5 for the refinement of macromolecular crystal structures. Acta Crystallographica Section D 67:355–367.

36. Emsley P, Cowtan K (2004) Coot: model-building tools for molecular graphics. Acta Crystallogr D Biol Crystallogr 60:2126–2132. PMID: 15572765 {Medline}

37. Cohen SX, Ben Jelloul M, Long F et al. (2008) ARP/wARP and molecular replacement: the next generation. Acta Crystallogr D Biol Crystallogr 64:49–60. PMID: 18094467 {Medline}

38. Painter J, Merritt EA (2006) TLSMD web server for the generation of multi-group TLS models. J Appl Cryst 39:109–111.

39. Chen VB, Arendall WB, 3rd, Headd JJ et al. (2010) MolProbity: all-atom structure validation for macromolecular crystallography. Acta Crystallogr D Biol Crystallogr 66:12–21. PMID: 20057044 {Medline}

40. Pettersen EF, Goddard TD, Huang CC et al. (2021) UCSF ChimeraX: Structure visualization for researchers, educators, and developers. Protein Sci 30:70–82. PMID: 32881101 {Medline}

41. Sievers F, Higgins DG (2021) The Clustal Omega Multiple Alignment Package. Methods Mol Biol 2231:3–16. PMID: 33289883 {Medline}

42. Robert X, Gouet P (2014) Deciphering key features in protein structures with the new ENDscript server. Nucleic Acids Res 42:W320–324. PMID: 24753421 {Medline}

